# *Ex-situ* mycoremediation of petroleum polluted soils in Ogoniland, Nigeria

**DOI:** 10.1101/2023.01.08.523087

**Authors:** Geoffrey O. Anoliefo, Martin Nwaokolo, Boniface O. Edegbai, Beckley Ikhajiagbe

## Abstract

The study investigated the mycoremediation oil-polluted soil from Ogoniland (at the B-dere/Nabem community).The mushroom, *Pleurotus tuberregium* was exposed to this soil as principal remediation component.While the fungus grew undisturbed, weeds were also allowed to grow from soil seed bank.Dominant weeds identified in the study include *Mariscus sp, Pepperomia pellucida, Synedrella nodiflora, Cyperus* sp., and *Oldenlandia corymbosa*. Survival and growth of microorganisms in the polluted soils were monitored.Bacterial species identified were *Escherichia coli, Micrococcus varians, Pseudomonas aeruginosa, P. vulgaris, Bacillus pumilis, Clostridum sp*., *Klebsiella*sp., and*Azotobacter* sp. The fungi species included *Mucormucedo, Aspergillus* sp., *Penicillin* sp., *Rhizopus* sp., *Trichoderma harzianum*, and *Fusarium solani*. The degradation of the petroleum pollutant wasmonitored through laboratory determination using the Gas Liquid Chromatographic methods. The determinations showed the gradual but steady decline in the concentration of the petroleum hydrocarbon pollutants in the Ogoni soil within the 4-month observation period. Significant reduction in heavy metal concentration of the soils was also evidenced.

## Introduction

Ogoni is a major ethnic group in the Niger Delta of Nigeria with an estimated population of about 750, 000 people (2006 Nigeria census). These people occupy a land area approximately 100, 000 km^2^on the South Eastern fringe of the Niger Delta. Ogoniland has 4Local Government Areas; Khana, Gokana, Tai and Eleme; and their traditional occupation are fishing with farming.Oil exploration activities are very high in Ogoniland given that crude oil was discovered in commercial quantity at Bomu in Gokana Local Government Area in 1958. Shell®currently has12 oil fields containing 116 drilled oil wells and 5 oil flow stations in Ogoniland. The petroleum and allied industries contribute a major portion of pollution in Nigeria. It is ironical that the Niger Delta with the highest deposit of crude oil and gas in the country has about the most fragile ecosystems in Nigeria. Most of the terrestrial ecosystems and shorelines in the oil-producing communities are important agricultural land under continuous cultivation. There is therefore need to provide adequate preventive measures to minimize the environmental pollution or intensify research activities at remediation strategies to mitigate the deleterious effects of oil on the ecosystem.

Oil in soil is quite damaging to resident plants and microbes. This is because oil polluted soils may become unsuitable due to a number of factors, including reduction in soil oxygen from the soil waxy nature, impenetrable nature and low infiltration, reduction in the level of available plant nutrients or a rise to a toxic level of heavy metals, as well as the presence of toxic aliphatic and aromatic compounds (Udo and Fayemi, 1975; Günther*et al*., 1996; Anoliefo et al., 2001; Anoliefo and Umweni 2004; Ikhajiagbe, 2010). Crude oil is physically, chemically and biologically harmful to soil because it contains many toxic compounds, such as polycyclic aromatic hydrocarbons, benzene and its substituted cycloalkane rings in relatively high concentrations and thus of serious concern worldwide (Barker and Gretchen 2002).These hydrocarbon compounds are also of high molecular weight with very low solubility in water thus preventing natural biodegradation process from working efficiently in hydrocarbon contaminated soils (Esin and Ayten, 2011). The compounds also penetrate macro-and micropores in soil and thus limit water and air transport that would be necessary for organic matter conversion.

There are many biological techniques used in the cleanup of land and water sources including bioventing, bio-slurping, hydraulic-pneumatic fracturing, soil bio-injection, air and water flushing, biopolymer shields, electro-bioreclamation and phytoremediation. However, most of these techniques are very expensive, preferred for cleanup of deep soil layers and may be limited in terms of soil properties and environmental conditions.

Bioremediation has become the most desirable approach to the cleanup of the environment. This is due to its low cost and ability to hinder the accumulation of contaminants (Anoliefo et al., 2010). Soil bioremediation is the process in which most of the organic pollutants are decomposed by soil microorganisms and converted to harmless products such as carbon dioxide, methane and water. No single microbial species is capable of degradation of all components of crude oil. Complete oil degradation requires simultaneous action of different microbial populations. Whereas plants have capacity environmental reclamation measures, soil microorganisms are a fundamentally important component of terrestrial habitats and their primary role govern the nutrient cycles and the maintenance of soil structure. In soil microbial realm, some microbes have the distinctive ability to degrade or convert organic pollutants to harmless biological products and so bioremediation mainly relies on the use of these microorganisms surviving in soil.

Plant may be used to increase the rates of hydrocarbon degradationby stimulating microbial growth and activity in the plant rhizosphere (Parrish *et al*., 2004). The rhizosphere is that zone of increased activity of microorganisms within the soil in very close proximity with the roots. Plangklang and Reungsang, (2008) reported that the soil in the rhizosphere soil generally consists of 10-100 times greater number of indigenous microorganisms than in bulk soil.Alexander (2000) reported that as contaminants age in soil, they undergo a process that limits bioavailability, where the chemicals move into soil micropores or soil organic matrix forming unextractible bound residues. Parrish *et al*.(2004) thus reported that fine plant roots are able to penetrate some of these pores, thereby increasing the contaminants available for degradation. Remediation activities usually rely on resident microorganisms as well as plant species. These plants emanate from soil seed banks. Kalamees and Zobel, (2002) reported that the recovery of vegetation after disturbance lie mainly in the buried seed populations.

Ogoniland has been in the ‘eye of the storm’, with respect to crude oil exploration and exploitation for more than four decades. This has inflicted hardship on the people, retarded development and marred individual self-actualization. Matters became worse when youths found alternative means of livelihood in the face of unemployment, deprivation and neglect. The vista of hope came in the form of pipeline vandalization through sabotage and breakage of crude oil delivery pipelines to siphon petroleum. The youths have since advanced their trade in the form construction of alternative non-licensed refineries. They conduct the refining of petroleum without safety checks and guidelines; thus giving way to incessant oil spills, leakages and accidents. For three years (2008 to 2011), an alternative (non-legalized) refinery was reportedly located around the boundary between B-dere and Nabem communities in Gokana L.G.A. in Rivers State. The non-licensed refining activity continued until a major pipeline was vandalised within the same area. The vandalized pipelines were allegedly promptly repaired but the spill had already occurred, and with the chronic spill from the alternative refineries, the community was almost over-run. The impacted area was reportedly within the B-dere community (Personal Communication). At the time of soil collection from the site, the contamination was very evident, as observed in and around River Muboo (Plate 1).

**Plate 1 (A - D).**
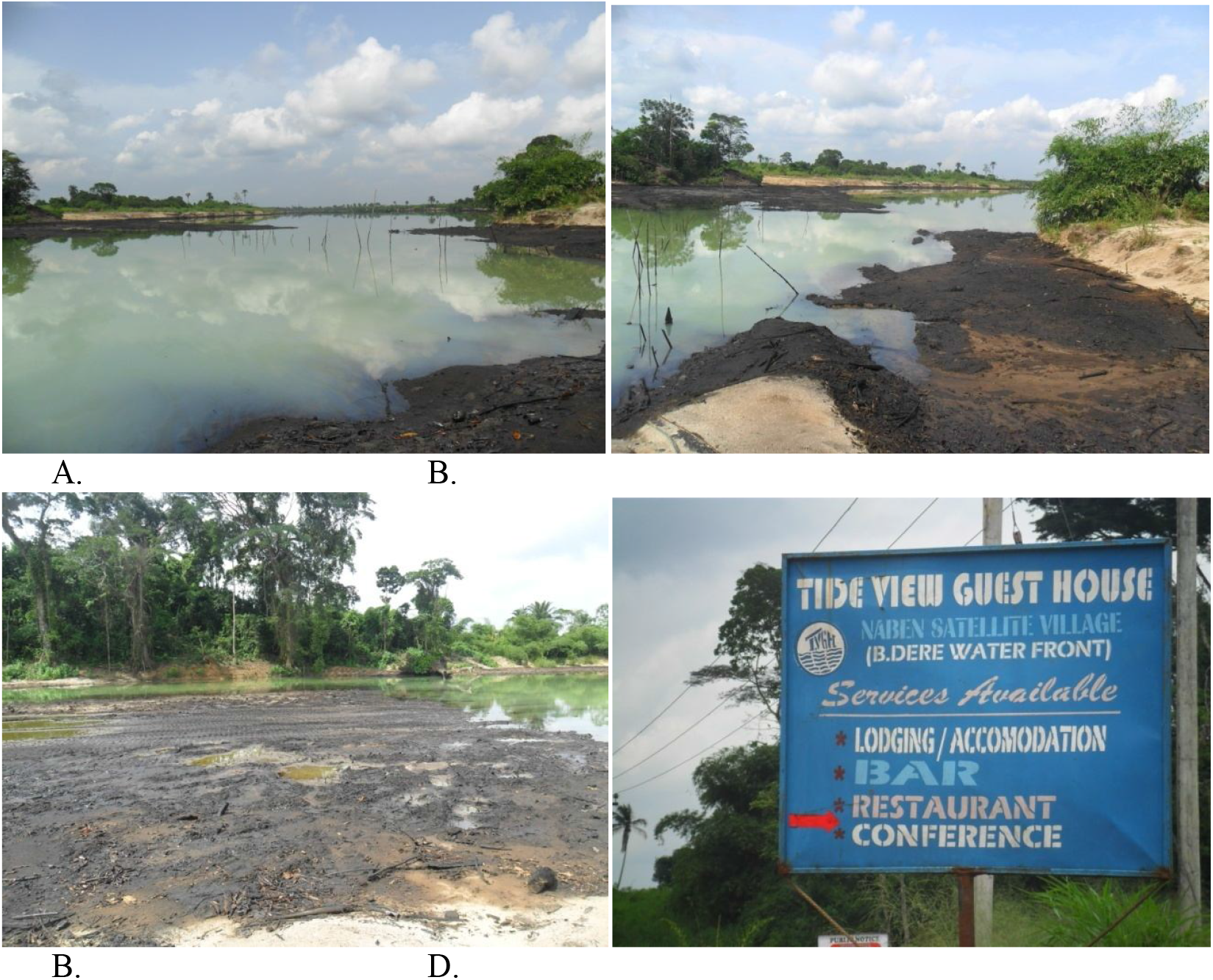
Site of soil sample collection within and around River Muboo.

The people ofOgoniland have largely remained agrarian and subsistent. Dependence on the government for regular clean up exercise, in cases of spills has not been successful. It was therefore necessary that the people be empowered to sometimes, attempt at cleaning with the available materials. The present study has utilized very basic, affordable and easily accessible materials for the remediation exercise. *Pleurotus tuberregium* is a common, edible mushroom that is easy to cultivate. The use of *Pleurotus tuberregium* was borne out of the consideration for the locals (farmers, fishermen and peasant traders) to conduct remediation activities using the prescribed materials. The sclerotia and sporophores are all edible and very well accepted by the local population. Cultivation of the mushroom requires agricultural wastes, which serve as substrates. The growth of the mushroom involves the degradation of the substrates and the pollutant in the soils. At the end of the exercise, the mushrooms are still very edible as the degradation is total (Anoliefo and Ikhajiagbe, 2012; Ikhajiagbe and Anoliefo, 2012a, b). The study would not involve any complex process and/or materials and as such can be learnt by the locals. Some attempts were made in 2012, by the Federal Ministry of Petroleum Resources, to clean up (Plate 2) the spill but until the period of soil collection, most of the physical presence of pollution was visible. The cleanup exercise probably led to the visible loss of vegetation in the area covered by the sign post (Plate 2).

**Plate 2.**
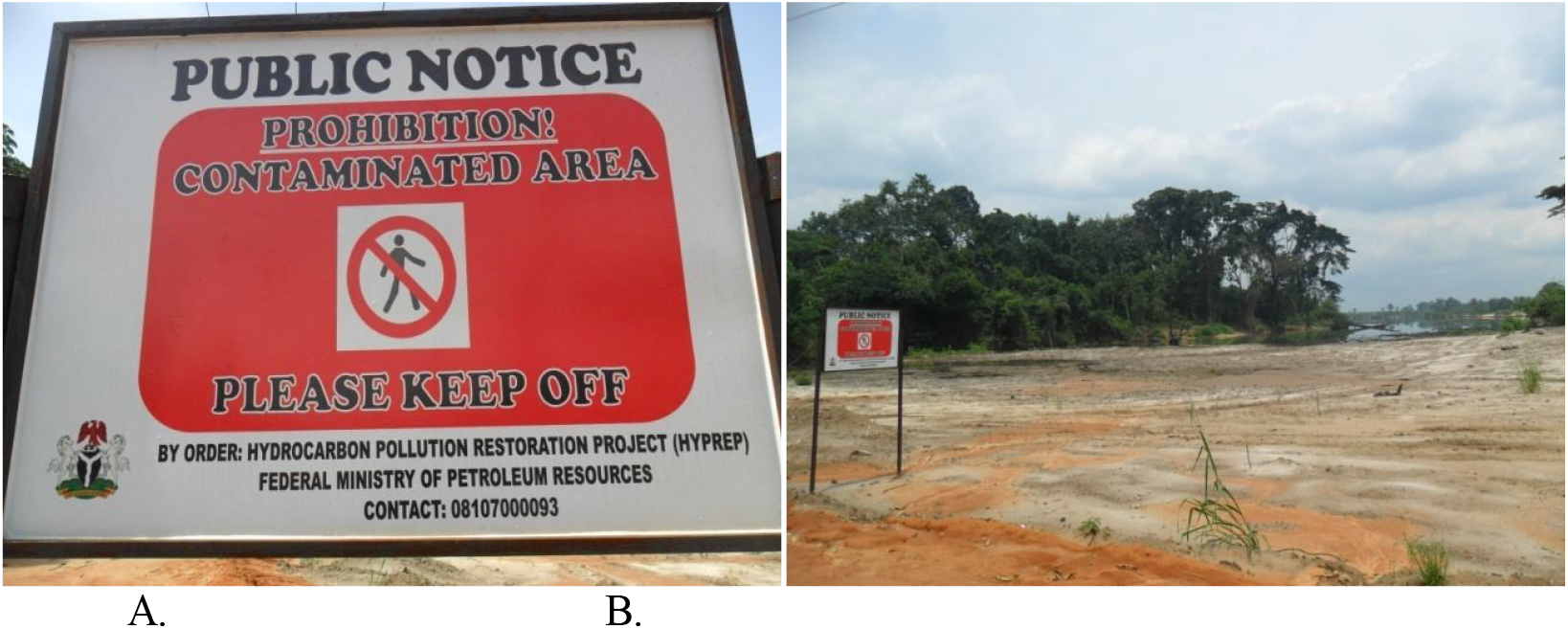
Evidence that cleanup activity had been carried out by Federal Ministry of Petroleum Resources.

Maintaining good soil quality and minimizing soil pollution remains essential in the realization of nature’s benefits from the land. A polluted soil would not be suitable for agriculture or other land uses. The aim of the study was to significantly degrade the petroleum pollutant using safe, biological means.The objectives of the study were to investigate the pivotal role attributed to the local mushroom in the remediation of the pollutants and to determine the extent to which synergism on the part of all the organismal actors in the remediation ‘theater’ was exhibited.

## Materials and Methods

### Collection of Ogoni soils and setting up of the experiment

The soils used in the study were obtained from randomly selected locations within the crude oil polluted sites in the B-dere and Nabem communities in Ogoniland, Rivers State. Soils were randomly collected from within.

The area delineated by the spill and upto 10 meters deep. The soils were later packed in four large sacs and transported to Benin City. The soils obtained were dark and oily, but were generally sandy. Care was taken to collect enough polluted soils for the study. Clean soil was obtained from an area that has never experienced pollution of any kind and used as control as well as diluent in combination with the polluted soils. Forty eight clean 25-liter buckets were purchased for dispensing of the soils.

### Preparation of Fungus

Healthy sclerotia of the mushroom, *Pleurotus tuberregium* used in the study were purchased from the Ikpoba Hill Market, Benin City.

### Soil Supplementation

Sawdust used as soil amendment agent in the study, was obtained from *Brachystegianigerica*, known for its ability to enhance growth and performance of the mushroom (Okhuoya*et al*., 1998). The sawdust was obtained from Ogida Sawmill in Benin City.

### Soil preparation

Soils collected from oil-polluted region in Ogoniland was herein referred to as stock soil (OG), whereas good soil (GS) referred to that obtained from a fallow portion of the Botanic Garden, University of Benin. In other to vary oil concentration in the soil, GS and OG were mixed to obtain 1, 5, 10, 25, 50 and 100 % concentrations. To obtain 1 % soil-soil mix, 1 part of OG was mixed with 99 parts of GS 99 parts of GS. The 100% concentration was pure OG. Each combination was then amended with 10 % (w/w) or 880 g of sawdust and mixed thoroughly.

### Seeding of Sclerotia

The whole experimental set up was allowed to naturally attenuate for 14 days following the methods of Ikhajiagbe et al. (2015). Thereafter sclerotia of the test fungus were obtained from Local Mushroom Growers in Benin City. An average of 880g (similar to 10% w/w of GS-OG-sawdust substrate) was required per experimental bucket. These were cut into smaller cubes and then seeded in each experimental bucket containing the GS-OG-sawdust mix (Table 1).

**Table 1:** Treatment options and coded identity in the study.

| **A.**<br>Concentration codes | Treatment Options | Replicates |
| --- | --- | --- |
| Control FO | Pure GS+sawdust (SD)+sclerotia (SC) | 3 |
| 1 % FO | 1 % GO/GS + SD+SC | 3 |
| 5 % FO | 5 % GO/GS +SD+SC | 3 |
| 10 % FO | 10 % GO/GS+SD+SC | 3 |
| 25 % FO | 25 % GO/GS+SD+SC | 3 |
| 50 % FO | 50 % GO/GS+SD+SC | 3 |
| 100 % FO | Pure GO+SC+SD | 3 |
| **B** |  |  |
| Control NO | Pure GS+SD only | 3 |
| 1 % NO | 1 % GO/GS+SD only | 3 |
| 5 % NO | 5 % GO/GS+SD only | 3 |
| 10 % NO | 10 % GO/GS+SD only | 3 |
| 25 % NO | 25 % GO/GS+SD only | 3 |
| 50 % NO | 50 % GO/GS+SD only | 3 |
| 100 % NO | 100 % GO/GS+SD only | 3 |
| GS | Pure GS only | 3 |
| GO | Pure GO only | 3 |

*NO=No fungi in Oil-polluted soil, FO= Fungi in Oil-polluted soil, GS= Good soil, GO= Oil-polluted soil, SD= Sawdust, SC= Sclerotia*

### Experimental Field Layout

The entire set up was left in an open shade for observation. The buckets were initially not perforated in order not to leach out the pollutant.

### Emergence of weeds from soil seed bank

After the harvest of mushrooms were harvested, the experimental set up was removed from the open shade and transferred into the open space. This was to allow for development of weeds from soil seed bank after soil remediation. The emerging weeds were identifed and then physically counted as long as each plant attained a minimum height of 5 cm.

### Microbial analysis

Resident soil microorganisms present in the control and experimental soils were monitored on interval basis through laboratory microbial analyses. Soil samples were collected from each of the replicate treatments and pooled together. The soil samples from each of the treatments and control were then air-dried, sieved, 1 g weight obtained and put into test tubes. Nine (9 ml) milliliter of normal saline solution was added to each test tube using a syringe and then stirred for 30 seconds with the Votex Genie mixer. The test tube with the soil in solution was covered with foil paper and then allowed to stay for 24 hours. After the waiting period (24 hours), serial dilution was carried out on the aliquot. This was done by further diluting the 1 ml of prepared aliquot with 9 ml of normal saline and stirring again for 30 seconds. The dilution was done for the second and third transfers to obtain a three-fold serial dilution.

### Media preparation

For the preparation of the medium for fungi culture, 39 g of potato dextrose agar (PDA) was weighed into conical flasks, mixed with 1 L of deionised water, autoclaved at a temperature of 121°C for 15 minutes at a pressure of 15 psi and then allowed to cool to 30°C. Two (2) capsules of chloramphenicol were added to inhibit the growth of bacteria. To culture for bacteria, 28 g of nutrient agar (NA) was weighed into conical flasks containing 1 L of deionised water, autoclaved for 15 minutes at a temperature of 121^0^C at 15 psi. Thereafter the medium was allowed to cool to 40°C. One (1) tablet of antibiotic (ketoconazole) was dissolved in the medium to inhibit the growth of fungi.

The working bench was kept sterile by swabbing with methylated spirit, Petri dishes used were labelled according to the treatment designation (1 to 100 % FO, 1 to 100 % NO, Control FO & NO, GS and OG). The medium which had cooled to body temperature was poured into different plates, (PDA for fungi and NA for bacteria) each having three (3) replicates. These activities were done under sterile environmental conditions and in addition, a lit burner was kept within the working bench to further maintain the sterile condition.

### Microbial colony count

Plates were divided into 4 quarters, the number of colonies in one quarter was counted and multiplied by four (4). For bacteria, colonies were counted from the base to ensure proper counting, while for the fungi, the colonies were counted from the top of the Petri dishes. Using these colony counts, the microbial load was calculated.

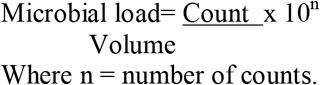

Where n = number of counts.

### Identification of bacteria and fungi species

Slides were prepared from the pure cultures. For bacteria identification, crystal violet, lugol iodine, ethyl alcohol and safranin were used. A flamed inoculating loop was used to pick organisms from the culture plate containing colonies and smeared on the slide. It was then passed through heat 4 times (heat fixing) and stained with the above stains respectively. Each stain was kept for two (2) minutes and rinsed with distilled water before staining with the next. For fungi, methyl blue was used to stain. After the slides were prepared, they were viewed under the Olympus binocular compound light microscope.

### Chemical analysis

#### Gas Chromatographic Studies

Soil samples were collected from control and treatments on the 31^st^ of July, 2014 for composite Total Petroleum Hydrocarbon (TPH) analysis. Extraction was carried out using the slightly modified procedure of Dean and Xiong (2000). Ten gram (10 g) of dry and homogenized soil sample was weighed, placed in an extraction thimble and then extracted using 200 ml of dichloromethane (DCM) via the reflux cycle for 15 hours for the extraction of polyaromatic hydrocarbons (PAH). Once the solvent was boiled, the vapour passed through a bypass arm into the condenser and condensed back onto the solvent in the thimble. As the solvent reached the top of the siphon arm, the solvent and extract were also siphoned back into the lower flask. The boiling/condensation cycle was repeated until all the samples were completely extracted into the lower flask.

Blanks were prepared following the same procedure without adding any soil sample. The standard sample used for quality control was prepared by adding the standard solution to DCM. All extracts were separated and activated copper was added to the combined extract for desulphurization. After subsequent drying over anhydrous sodium sulphate, the extract was concentrated to 1.0 ml using the rotary evaporator. An internal standard mixture (naphthalene-d8, acenaphthene-d10, phenanthrene-d10, chrysene-d12, and perylene-d12) solution was then added to the extract for analysis using the Hewlett Packard HP 5890 series II Gas Chromatograph with mass selective detection.

#### GC-MS Instrumentation and Conditions

The Hewlett Packard HP 5890 series II Gas Chromatograph, equipped with an Agilent 7683B Injector (Agilent Technologies, Santa Clara, CA, USA), a 30 m, 0.25 mm i.d. HP-5MS capillary column (Hewlett-Packard, Palo Alto, CA, USA) coated with 5% phenyl-methylsiloxane (film thickness 0.25 µm) and an Agilent 5975 mass selective detector (MSD) was used to separate and quantify the PAH compounds. The samples were injected in the splitless mode at an injection temperature of 300 °C. The transfer line and ion source temperatures were 280 °C and 200 °C. The column temperature was initially held at 40 °C for 1 min, raised to 120 °C at the rate of 25 °C per minute, then to 160 °C at the rate of 10 °C per minute, and finally to 300 °C at the rate of 5 °C per minute, held at final temperature for 15 minutes. The detector temperature was kept at 280 °C. Helium was used as a carrier gas at a constant flow rate of 1 mL per minute. Mass spectrometry was acquired using the electron ionization (EI) and selective ion monitoring (SIM) modes.

#### Determination of heavy metals in soil using the Atomic Absorption Spectrophotometer (AAS) and employing the Wet Digestion method

This method is for the rapid determination of Cu, Zn, As, Pb, Cr, V, Ni, Cd, Fe and Mn, in soil samples using AAS after double acid extraction. In the technique, the soil samples were not completely digested. However, the labile fractions of the metals were leached into the extracted solution. The samples were placed in glass Petri dishes and sun dried for 24 hours. After 24 hours of drying, lumps present were broken up with a clean glass rod in order to expose the inside for drying.When the samples appeared to be dried, they were left under the sun for further 24 hours before grinding.After drying, the soil was ground. In heavy contaminated soil, it was necessary to break up the hard pieces using a mortar and pestle.

##### Extraction proceedure

One gram (1 g) of the dried soil sample was transfered into an acid–wash 250 ml extraction bottle.Nine (9) ml of concentrated HCl, 3ml of HNO_3_ and 2ml of perchloric acid were added. The mixture was digested for 6 hours on the mechanical-shaker-hot-plate. After digestion was completed, 20 ml of distilled water was added and the solution was filtered through a whatman No 42 filter paper and finally made up to 100 ml mark. Blank samples were **p**repared.The filtrate was analyzed for metals using the Atomic Absorption PG 550 Spectrometer.

### Calibration and analysis

Single elemental standards were prepared by diluting 1000 mg/L stock solutions of the respective individual elements (Cu, As, Zn, Pb, Cr, V, Ni, Cd, Fe, Mn). A minimum of five standard working solutions were prepared daily from the stock Solutions ranged from 0.1mg/l to 1 mg/l.External calibration was used by running deionised water and a suite of calibration standards for each element, and calibration curve was then generated for each individual metal.The extracted solutions and blank were then run on the AA to obtain the absorbance values, and the concentrations of each metal in the digested samples were automatically calculated from the equation of the calibration curve by the AAS equipment.

For quality assurance, field acidified deionised water was first aspirated as blanks in duplicates and laboratory control samples were run as Quality Control samples.

### Statistics

Means of data were calculated and separated by using the Least Significant Difference. Other forms of statistics were those of ecological significance that required comparism with standard benchmark (Efroymson*et al*., 1997a, b; Cal-EPA, 2005).

## Results

Results showed that there were basically two prominent weeds in the oil-polluted soil - *Mariscus* sp., *Synedrella nodiflora*. There were more weeds in the NO-soil compared to when soil was amended with sawdust and then inoculated with test mushroom (Table 2).Heavy Metal composition content of soil two weeks after amendment with sawdust, before introduction of Sclerotia have been presented on Table 3a. Compared to soil without fungus, the fungus amended soil had less contamination with heavy metals after the study period indicating better remediation potentials than when soil was not augmented with mushroom. Fe concentration was 10.02mg/kg in FO(100%)-soil compared to 59.74 mg/kg in NO(100%)-soil. Similarly, concentration of arsenic was non-detectable in FO-soils, but was detected in NO-soils at 2 weeks after exposure to test fungus. However at 4 months, remediation capacities were comparable (Table 3b).

**Table 2:** Growth of weeds from soil seed bank in control and treated soils in 60 days of observation

| Treatment | Weeds observed in FO soils with species number | Total |
| --- | --- | --- |
| Good Soil | *Mariscus* sp (20), *Pepperomia pellucida*(10), *Synedrella nodiflora*(25) and *Oldenlandia corymbosa*(05) | 60 |
| Control FO | *Mariscus* sp.(07), *Synedrella nodiflora*(09) | 16 |
| 1 % FO | *Mariscus* sp.(14), *Synedrella nodiflora*(24) | 38 |
| 5 % FO | *Mariscus* sp.(35), *Synedrella nodiflora*(5), *Oldenlandia corymbosa*(1) | 41 |
| 10 % FO | *Mariscus* sp.(9), *Synedrella nodiflora*(33) | 42 |
| 25 % FO | *Mariscus* sp.(13), *Synedrella nodiflora*(11) | 24 |
| 50 % FO | *Mariscus* sp.(05), *Pepperomia pellucida*(07) | 12 |
| 100 % FO | No weed was observed | 0 |
| 1 % NO | *Mariscus* sp.(28), *Synedrella nodiflora*(52) | 80 |
| 5 % NO | *Mariscus* sp.(54), *Synedrella nodiflora*(40). | 94 |
| 10 % NO | *Mariscus* sp.(34), *Synedrella nodiflora*(32) | 68 |
| 25 % NO | *Mariscus* sp.(18), *Synedrella nodiflora*(32), *Pepperomia pellucida* (21) | 71 |
| 50 % NO | *Synedrella nodiflora* (9), *Mariscus* sp.(05), *Drymaria cordata*(10) | 52 |
| 100 % NO | No weed was observed | 0 |

**Table 3:**
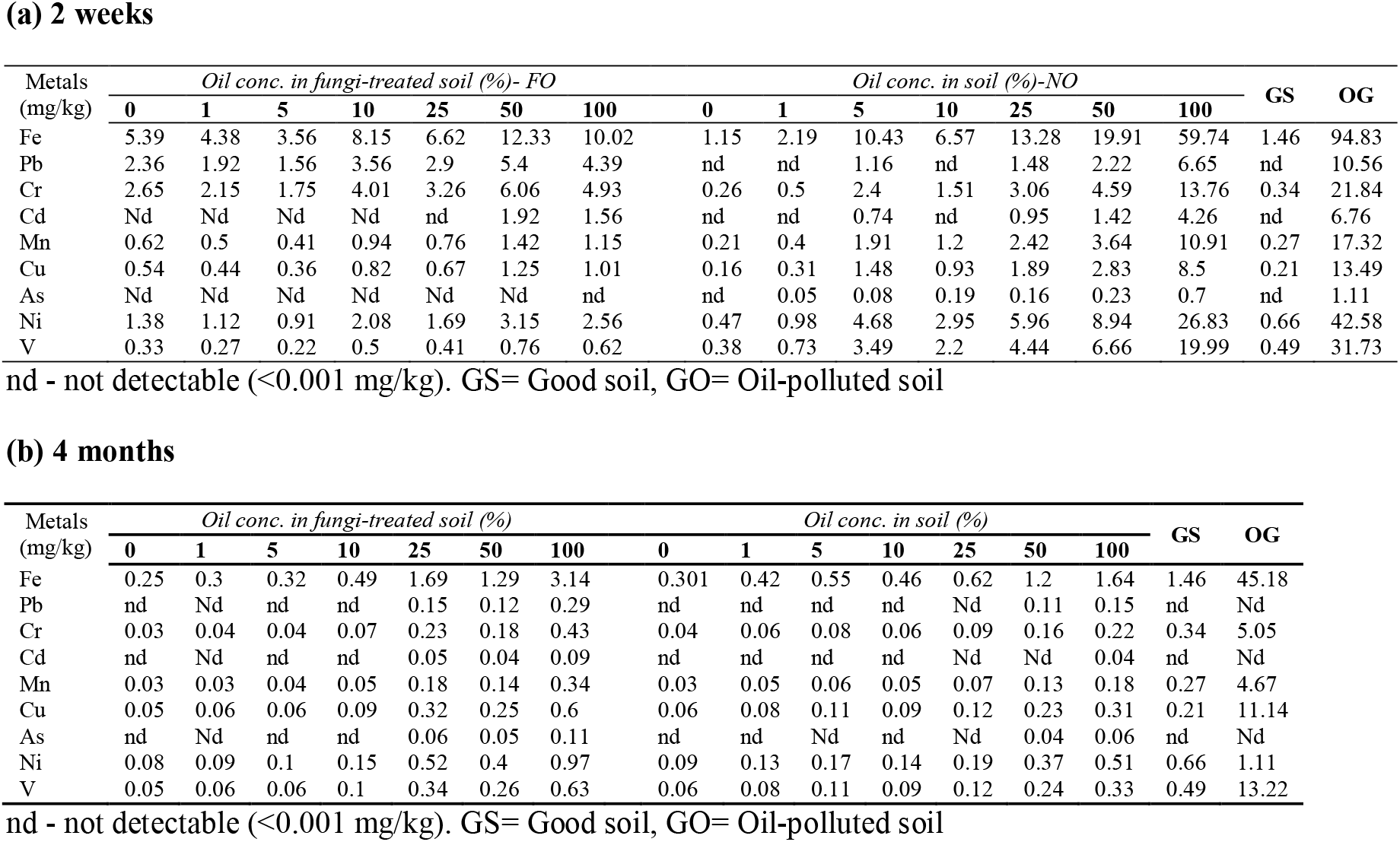
Heavy Metal composition content of soilafter amendment with sawdust, before introduction of Sclerotia

**(a) 2 weeks**
| Metals<br>(mg/kg) | *Oil conc. in fungi-treated soil (%) - FO* |  |  |  |  |  |  | *Oil conc. in soil (%) - NO* |  |  |  |  |  |  | GS | OG |
| --- | --- | --- | --- | --- | --- | --- | --- | --- | --- | --- | --- | --- | --- | --- | --- | --- |
|  | 0 | 1 | 5 | 10 | 25 | 50 | 100 | 0 | 1 | 5 | 10 | 25 | 50 | 100 |  |  |
| Fe | 5.39 | 4.38 | 3.56 | 8.15 | 6.62 | 12.33 | 10.02 | 1.15 | 2.19 | 10.43 | 6.57 | 13.28 | 19.91 | 59.74 | 1.46 | 94.83 |
| Pb | 2.36 | 1.92 | 1.56 | 3.56 | 2.9 | 5.4 | 4.39 | nd | nd | 1.16 | nd | 1.48 | 2.22 | 6.65 | nd | 10.56 |
| Cr | 2.65 | 2.15 | 1.75 | 4.01 | 3.26 | 6.06 | 4.93 | 0.26 | 0.5 | 2.4 | 1.51 | 3.06 | 4.59 | 13.76 | 0.34 | 21.84 |
| Cd | Nd | Nd | Nd | Nd | nd | 1.92 | 1.56 | nd | nd | 0.74 | nd | 0.95 | 1.42 | 4.26 | nd | 6.76 |
| Mn | 0.62 | 0.5 | 0.41 | 0.94 | 0.76 | 1.42 | 1.15 | 0.21 | 0.4 | 1.91 | 1.2 | 2.42 | 3.64 | 10.91 | 0.27 | 17.32 |
| Cu | 0.54 | 0.44 | 0.36 | 0.82 | 0.67 | 1.25 | 1.01 | 0.16 | 0.31 | 1.48 | 0.93 | 1.89 | 2.83 | 8.5 | 0.21 | 13.49 |
| As | Nd | Nd | Nd | Nd | Nd | Nd | nd | nd | 0.05 | 0.08 | 0.19 | 0.16 | 0.23 | 0.7 | nd | 1.11 |
| Ni | 1.38 | 1.12 | 0.91 | 2.08 | 1.69 | 3.15 | 2.56 | 0.47 | 0.98 | 4.68 | 2.95 | 5.96 | 8.94 | 26.83 | 0.66 | 42.58 |
| V | 0.33 | 0.27 | 0.22 | 0.5 | 0.41 | 0.76 | 0.62 | 0.38 | 0.73 | 3.49 | 2.2 | 4.44 | 6.66 | 19.99 | 0.49 | 31.73 |

**(b) 4 months**
| Metals<br>(mg/kg) | *Oil conc. in fungi-treated soil (%)* |  |  |  |  |  |  | *Oil conc. in soil (%)* |  |  |  |  |  |  | GS | OG |
| --- | --- | --- | --- | --- | --- | --- | --- | --- | --- | --- | --- | --- | --- | --- | --- | --- |
|  | 0 | 1 | 5 | 10 | 25 | 50 | 100 | 0 | 1 | 5 | 10 | 25 | 50 | 100 |  |  |
| Fe | 0.25 | 0.3 | 0.32 | 0.49 | 1.69 | 1.29 | 3.14 | 0.301 | 0.42 | 0.55 | 0.46 | 0.62 | 1.2 | 1.64 | 1.46 | 45.18 |
| Pb | nd | Nd | nd | nd | 0.15 | 0.12 | 0.29 | nd | nd | nd | nd | Nd | 0.11 | 0.15 | nd | Nd |
| Cr | 0.03 | 0.04 | 0.04 | 0.07 | 0.23 | 0.18 | 0.43 | 0.04 | 0.06 | 0.08 | 0.06 | 0.09 | 0.16 | 0.22 | 0.34 | 5.05 |
| Cd | nd | Nd | nd | nd | 0.05 | 0.04 | 0.09 | nd | nd | nd | nd | Nd | Nd | 0.04 | nd | Nd |
| Mn | 0.03 | 0.03 | 0.04 | 0.05 | 0.18 | 0.14 | 0.34 | 0.03 | 0.05 | 0.06 | 0.05 | 0.07 | 0.13 | 0.18 | 0.27 | 4.67 |
| Cu | 0.05 | 0.06 | 0.06 | 0.09 | 0.32 | 0.25 | 0.6 | 0.06 | 0.08 | 0.11 | 0.09 | 0.12 | 0.23 | 0.31 | 0.21 | 11.14 |
| As | nd | Nd | nd | nd | 0.06 | 0.05 | 0.11 | nd | nd | Nd | nd | Nd | 0.04 | 0.06 | nd | Nd |
| Ni | 0.08 | 0.09 | 0.1 | 0.15 | 0.52 | 0.4 | 0.97 | 0.09 | 0.13 | 0.17 | 0.14 | 0.19 | 0.37 | 0.51 | 0.66 | 1.11 |
| V | 0.05 | 0.06 | 0.06 | 0.1 | 0.34 | 0.26 | 0.63 | 0.06 | 0.08 | 0.11 | 0.09 | 0.12 | 0.24 | 0.33 | 0.49 | 13.22 |

nd - not detectable (<0.001 mg/kg). GS= Good soil, GO= Oil-polluted soil

Heavy metal content in mushroom fruiting body was dose-dependent. At 50% concentration of polluted soil, Cd was undetected (<0.001 mg/kg) in the mushroom fruiting body. Irrespective of soil concentration, V was not detected in fruiting body (Table 4)

**Table 4:** Heavy Metal content of mushroom fruiting body harvested from a crude oil-polluted soil amendment with the mushroom sclerotia for 4 months

| Metals | *Oil conc. in fungi-treated soil (%)* |  |  |  |  |  |  |
| --- | --- | --- | --- | --- | --- | --- | --- |
|  | 0 | 1 | 5 | 10 | 25 | 50 | 100 |
| Fe | 0.2 | 1.47 | 1.26 | 1.68 | 4.33 | 1.89 | 6.03 |
| Pb | 0.02 | 0.14 | 0.12 | 0.16 | 0.42 | 0.18 | 1.00 |
| Cr | 0.05 | 0.37 | 0.31 | 0.42 | 1.08 | 0.47 | 2.57 |
| Cd | nd | nd | Nd | nd | nd | 0.01 | 0.03 |
| Mn | 0.05 | 0.38 | 0.32 | 0.43 | 1.11 | 0.49 | 2.65 |
| Cu | 0.07 | 0.51 | 0.44 | 0.59 | 0.15 | 0.66 | 0.36 |
| As | 0.04 | 0.29 | 0.25 | 0.33 | 0.86 | 0.38 | 0.21 |
| Ni | 0.11 | 0.8 | 0.69 | 0.92 | 0.24 | 1.03 | 0.56 |
| V | nd | nd | Nd | nd | nd | nd | nd |

nd - not detectable (<0.001 mg/kg).

Table 5 presents the bioaccumulation factors (BF) for heavy metals accumulated in the fruiting bodies of *P. tuberregium* after harvest from the crude oil-polluted soil two months after. Significant accumulations of heavy metals in the fruiting bodies are evident when BF > 1. The most significant accumulation of heavy metal ws seen in Cd (300 units). Least accumulations in the fruiting bodies was recorded for V, irrespective of concentration in soil (Table 5). In the sclerotia however, higher depositions of V were recorded (1.18 < BF < 1.82 units).

**Table5:** Bioaccumulation factors (BF) for heavy metals accumulated in fruiting after harvest from a crude oil-polluted soil two months after it was buried for remediation purpose

| Metals | Oil conc. in fungi-treated soil (%) |  |  |  |  |  |
| --- | --- | --- | --- | --- | --- | --- |
|  | 1 | 5 | 10 | 25 | 50 | 100 |
| **Fruiting bodies** |  |  |  |  |  |  |
| Fe (0.200) | 7.35 | 6.30 | 8.40 | 21.65 | 9.45 | 30.15 |
| Pb (0.020) | 7.00 | 6.00 | 8.00 | 21.00 | 9.00 | 50.00 |
| Cr (0.050) | 7.40 | 6.20 | 8.40 | 21.60 | 9.40 | 51.40 |
| Cd (<0.001) | 1.00 | 1.00 | 1.00 | 1.00 | 100.00 | 300.00 |
| Mn (0.050) | 7.60 | 6.40 | 8.60 | 22.20 | 9.80 | 53.00 |
| Cu (0.070) | 7.29 | 6.29 | 8.43 | 2.14 | 9.43 | 5.14 |
| As (0.040) | 7.25 | 6.25 | 8.25 | 21.50 | 9.50 | 5.25 |
| Ni (0.110) | 7.27 | 6.27 | 8.36 | 2.18 | 9.36 | 5.09 |
| V (<0.001) | 1.00 | 1.00 | 1.00 | 1.00 | 1.00 | 1.00 |
| **Sclerotia** |  |  |  |  |  |  |
| Fe (1.130) | 1.16 | 0.13 | 1.04 | 1.61 | 6.97 | 7.74 |
| Pb (<0.001) | 1.00 | 1.00 | 1.00 | 1.00 | 1.00 | 1.00 |
| Cr (0.320) | 1.19 | 0.56 | 1.50 | 1.63 | 16.88 | 10.38 |
| Cd (<0.001) | 1.00 | 1.00 | 1.00 | 1.00 | 1.00 | 1.00 |
| Mn (1.800) | 0.12 | 0.13 | 0.15 | 0.16 | 1.67 | 1.76 |
| Cu (0.670) | 1.18 | 1.34 | 1.51 | 1.67 | 1.67 | 1.78 |
| As (<0.001) | 1.00 | 1.00 | 1.00 | 1.00 | 1.00 | 1.00 |
| Ni (0.750) | 1.19 | 0.13 | 1.51 | 1.63 | 1.68 | 6.97 |
| V (0.220) | 1.18 | 1.36 | 1.55 | 1.68 | 1.68 | 1.82 |

BF > 1 indicates significant heavy metal bioaccumulation. Values in parentheses are reference values; these are values obtained in fruiting bodies and sclerotia sown in non-polluted soil/control

Percentage reduction in TPH at 2 weeks was higher in the fungus-amended soil (FO), compared to NO when pollutant concentrations were higher. There was a 95.21% reduction in TPH in 100%FO compared to 11.39% in 100%NO (Table 6). However, at 4 months, reduction percentage was comparative (97.51 – 97.99%). The implication is that reduction in TPH took shorter time in the fungus-amended soil compared to when so was not amended.

**Table 6:**
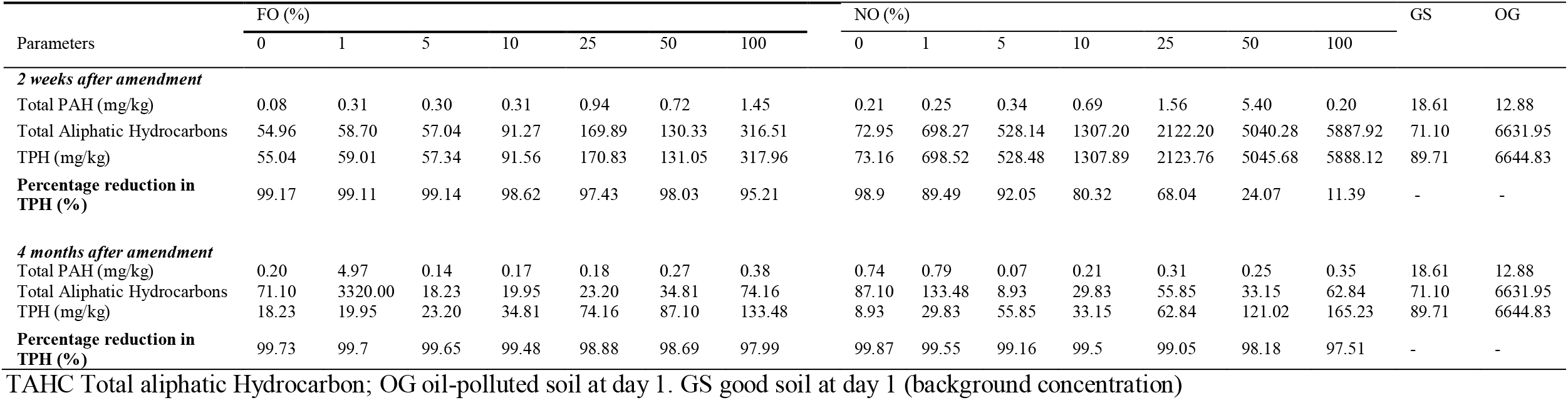
Total Petroleum Hydrocarbon content of soil after amendment with sawdust, before introduction of Sclerotia

| Parameters | FO (%) |  |  |  |  |  |  | NO (%) |  |  |  |  |  |  | GS | OG |
| --- | --- | --- | --- | --- | --- | --- | --- | --- | --- | --- | --- | --- | --- | --- | --- | --- |
|  | 0 | 1 | 5 | 10 | 25 | 50 | 100 | 0 | 1 | 5 | 10 | 25 | 50 | 100 |  |  |
| 2 weeks after amendment |  |  |  |  |  |  |  |  |  |  |  |  |  |  |  |  |
| Total PAH (mg/kg) | 0.08 | 0.31 | 0.30 | 0.31 | 0.94 | 0.72 | 1.45 | 0.21 | 0.25 | 0.34 | 0.69 | 1.56 | 5.40 | 0.20 | 18.61 | 12.88 |
| Total Aliphatic Hydrocarbons | 54.96 | 58.70 | 57.04 | 91.27 | 169.89 | 130.33 | 316.51 | 72.95 | 698.27 | 528.14 | 1307.20 | 2122.20 | 5040.28 | 5887.92 | 71.10 | 6631.95 |
| TPH (mg/kg) | 55.04 | 59.01 | 57.34 | 91.56 | 170.83 | 131.05 | 317.96 | 73.16 | 698.52 | 528.48 | 1307.89 | 2123.76 | 5045.68 | 5888.12 | 89.71 | 6644.83 |
| Percentage reduction in TPH (%) | 99.17 | 99.11 | 99.14 | 98.62 | 97.43 | 98.03 | 95.21 | 98.9 | 89.49 | 92.05 | 80.32 | 68.04 | 24.07 | 11.39 | - | - |
| 4 months after amendment |  |  |  |  |  |  |  |  |  |  |  |  |  |  |  |  |
| Total PAH (mg/kg) | 0.20 | 4.97 | 0.14 | 0.17 | 0.18 | 0.27 | 0.38 | 0.74 | 0.79 | 0.07 | 0.21 | 0.31 | 0.25 | 0.35 | 18.61 | 12.88 |
| Total Aliphatic Hydrocarbons | 71.10 | 3320.00 | 18.23 | 19.95 | 23.20 | 34.81 | 74.16 | 87.10 | 133.48 | 8.93 | 29.83 | 55.85 | 33.15 | 62.84 | 71.10 | 6631.95 |
| TPH (mg/kg) | 18.23 | 19.95 | 23.20 | 34.81 | 74.16 | 87.10 | 133.48 | 8.93 | 29.83 | 55.85 | 33.15 | 62.84 | 121.02 | 165.23 | 89.71 | 6644.83 |
| Percentage reduction in TPH (%) | 99.73 | 99.7 | 99.65 | 99.48 | 98.88 | 98.69 | 97.99 | 99.87 | 99.55 | 99.16 | 99.5 | 99.05 | 98.18 | 97.51 | - | - |

TAHC Total aliphatic Hydrocarbon; OG oil-polluted soil at day 1. GS good soil at day 1 (background concentration)

Four months after exposure to fungus, Polycyclic Aromatic Hydrocarbon (PAH) content of soil in the fungus-amended soil, comparative to that which was not amended, was comparable (Table 7). Concentrations were significantly low compared to original pollution levels (OG). Benz[a]anthracene, Benzo[k]fluoranthene andChrysene were undetected after soil was amended with fungus. There has been over 95% reduction in total aliphatic hydrocarbons in the fungus-amended oil polluted soil compared to background concentration in Ogoni soil (Table 8).

**Table 7:** Polycyclic Aromatic Hydrocarbon content of soil 4 months after amendment with sawdust, before introduction of Sclerotia

| Parameters | FO (%) |  |  |  |  |  |  | NO (%) |  |  |  |  |  |  | GS4m | OG4m |
| --- | --- | --- | --- | --- | --- | --- | --- | --- | --- | --- | --- | --- | --- | --- | --- | --- |
|  | 0 | 1 | 5 | 10 | 25 | 50 | 100% | 0 | 1 | 5 | 10 | 25 | 50 | 100 |  |  |
| Naphthalene (NA) | 0.017 | 0.02 | 0.021 | 0.032 | 0.135 | 0.087 | 0.264 | 0.008 | 0.025 | 0.036 | 0.03 | 0.041 | 0.079 | 0.107 | 0.103 | 2.698 |
| Acenaphthene (AC) | 0.011 | 0.013 | 0.013 | 0.02 | 0.028 | 0.055 | 0.105 | 0.005 | 0.016 | 0.023 | 0.019 | 0.026 | 0.05 | 0.069 | 0.011 | 1.7493 |
| Acenaphthylene (ACN) | 0.009 | 0.011 | 0.012 | 0.018 | 0.021 | 0.048 | 0.078 | 0.005 | 0.014 | 0.02 | 0.017 | 0.023 | 0.044 | 0.06 | <0.001 | 0.1113 |
| Fluorene (FL) | 0.003 | 0.004 | 0.004 | 0.006 | 0.152 | 0.016 | 0.021 | 0.002 | 0.005 | 0.007 | 0.006 | 0.008 | 0.015 | 0.02 | <0.001 | 0.6111 |
| Phenanthrene (PHE) | 0.002 | 0.002 | 0.002 | 0.004 | <0.001 | 0.01 | 0.008 | <0.001 | 0.003 | 0.004 | <0.001 | 0.005 | 0.009 | 0.012 | 0.052 | 0.0147 |
| Anthracene (AN) | 0.001 | 0.001 | 0.001 | 0.002 | 0.021 | 0.005 | 0.013 | <0.001 | 0.001 | 0.002 | <0.001 | 0.002 | 0.004 | 0.006 | <0.001 | 2.1441 |
| Fluoranthene (FA) | 0.001 | 0.001 | 0.001 | 0.001 | <0.001 | 0.003 | 0.008 | <0.001 | 0.001 | 0.001 | <0.001 | 0.001 | 0.003 | 0.004 | <0.001 | 2.1042 |
| Pyrene (PY) | 0.001 | 0.001 | 0.001 | 0.002 | <0.001 | 0.005 | 0.048 | <0.001 | 0.001 | 0.002 | <0.001 | 0.002 | 0.004 | 0.006 | 0.003 | 1.021 |
| Benz[a]anthracene (B[a]A) | <0.001 | 0.001 | 0.001 | 0.001 | 0.011 | 0.002 | 0.025 | <0.001 | 0.001 | 0.001 | <0.001 | 0.001 | 0.002 | 0.003 | <0.001 | 1.002 |
| Chrysene (CHR) | <0.001 | <0.001 | <0.001 | <0.001 | <0.001 | <0.001 | <0.001 | <0.001 | <0.001 | <0.001 | <0.001 | <0.001 | <0.001 | <0.001 | <0.001 | 0.039 |
| Benzo[b]fluoranthene (B[b]F) | 0.033 | 0.039 | 0.041 | 0.062 | <0.001 | 0.17 | 0.063 | 0.016 | 0.049 | 0.071 | 0.059 | 0.08 | 0.154 | 0.21 | 0.032 | 1.9234 |
| Benzo[k]fluoranthene (B[k]F) | <0.001 | <0.001 | <0.001 | <0.001 | <0.001 | <0.001 | <0.001 | <0.001 | <0.001 | <0.001 | <0.001 | <0.001 | <0.001 | <0.001 | <0.001 | 2.931 |
| Benzo[a]pyrene (B[a]P) | 0.027 | 0.032 | 0.034 | 0.051 | 0.014 | 0.139 | 0.157 | 0.013 | 0.04 | 0.058 | 0.048 | 0.065 | 0.126 | 0.172 | <0.001 | 1.44 |
| Indeno[1, 2, 3-cd]-pyrene (IP) | 0.038 | 0.044 | 0.048 | 0.072 | <0.001 | 0.196 | 0.04 | 0.019 | 0.056 | 0.082 | 0.068 | 0.092 | 0.177 | 0.242 | <0.001 | 0.824 |
| **Total PAH (mg/kg)** | 0.141 | 0.167 | 0.179 | 0.269 | 0.382 | 0.736 | 0.79 | 0.069 | 0.21 | 0.308 | 0.248 | 0.346 | 0.667 | 0.91 | 0.201 | 18.613 |

TAHC Total aliphatic Hydrocarbon; OG4m oil-polluted soil at 4 months. GS4m good soil at 4 months

**Table 8:** Total Aliphatic Hydrocarbon content of soil at 4 monthsafter amendment with sawdust, before introduction of Sclerotia

| Parameters | FO (%) |  |  |  |  |  |  | NO (%) |  |  |  |  |  |  | GS | OG |
| --- | --- | --- | --- | --- | --- | --- | --- | --- | --- | --- | --- | --- | --- | --- | --- | --- |
|  | 0 | 1 | 5 | 10 | 25 | 50 | 100% | 0 | 1 | 5 | 10 | 25 | 50 | 100 |  |  |
| Nonane (C 9) | 8.880 | 10.490 | 11.300 | 16.950 | 29.150 | 42.790 | 51.310 | 10.490 | 14.530 | 19.370 | 16.150 | 21.800 | 41.980 | 57.320 | 35.880 | 2242.620 |
| Decane(C 10) | 4.850 | 5.730 | 6.170 | 9.250 | 13.960 | 13.750 | 28.010 | 2.200 | 7.930 | 10.580 | 8.810 | 11.900 | 22.910 | 31.290 | 8.590 | 1073.960 |
| Dodecane(C 12) | 2.220 | 2.620 | 2.820 | 4.240 | 6.810 | 6.700 | 12.820 | 0.210 | 3.630 | 4.840 | 4.030 | 5.450 | 10.490 | 14.320 | 5.760 | 523.720 |
| Tetradecane(C 14) | 2.290 | 1.100 | 2.910 | 4.360 | 9.500 | 9.850 | 13.210 | 0.020 | 3.740 | 4.990 | 4.160 | 5.610 | 10.810 | 14.750 | 4.320 | 730.580 |
| Hexadecane(C 16) | <0.001 | <0.001 | <0.001 | <0.001 | 0.000 | 0.000 | 0.000 | <0.001 | <0.001 | 0.050 | <0.001 | 0.060 | 0.110 | 0.150 | 0.720 | 45.260 |
| Octadecane(C 18) | <0.001 | <0.001 | <0.001 | <0.001 | 8.930 | 8.790 | 17.920 | <0.001 | <0.001 | 6.770 | <0.001 | 7.610 | 14.660 | 20.010 | 3.680 | 687.020 |
| Nonadecane(C 19) | <0.001 | <0.001 | <0.001 | <0.001 | 0.000 | 0.000 | 0.000 | <0.001 | <0.001 | 2.610 | <0.001 | 2.930 | 5.650 | 7.720 | 2.850 | 395.850 |
| Eicosane(C 20) | <0.001 | <0.001 | <0.001 | <0.001 | 0.000 | 0.000 | 0.000 | <0.001 | <0.001 | 1.880 | <0.001 | 2.120 | 4.080 | 5.570 | 2.720 | 264.120 |
| Docasane(C 22) | <0.001 | <0.001 | <0.001 | <0.001 | 5.800 | 5.710 | 10.210 | <0.001 | <0.001 | 3.860 | <0.001 | 4.340 | 8.350 | 11.410 | 1.740 | 446.550 |
| Tetracosane(C 24) | <0.001 | <0.001 | <0.001 | <0.001 | 0.000 | 0.000 | 0.000 | <0.001 | <0.001 | 0.580 | <0.001 | 0.660 | 1.260 | 1.730 | 1.240 | 108.860 |
| hexacosane(C 26) | <0.001 | <0.001 | <0.001 | <0.001 | 0.000 | 0.000 | 0.000 | <0.001 | <0.001 | 0.260 | <0.001 | 0.290 | 0.560 | 0.770 | 0.630 | 65.630 |
| Tricosane(C 30) | <0.001 | <0.001 | <0.001 | <0.001 | 0.000 | 0.000 | 0.000 | <0.001 | <0.001 | 0.070 | <0.001 | 0.070 | 0.140 | 0.190 | 2.720 | 60.410 |
| TAHC | 18.230 | 19.950 | 23.200 | 34.810 | 74.160 | 87.100 | 133.480 | 8.930 | 29.830 | 55.850 | 33.150 | 62.840 | 121.020 | 165.230 | **71.100** | 6644.830 |

TAHC Total aliphatic Hydrocarbon; OG oil-polluted soil at day 1. GS good soil at day 1

PAH content of sclerotia after harvest from oil-polluted soil/substrate at 4 months after exposure showed that PAH contents were either negligible or not detected at all (Table 9).

**Table 9:** Polycyclic Aromatic Hydrocarbon content of sclerotia after it was harvested from an oil-polluted soil 4 months after it was buried for remediation purpose

| Parameters | 0% FO | 1% FO | 5% FO | 10% FO | 25% FO | 50% FO | 100% FO |
| --- | --- | --- | --- | --- | --- | --- | --- |
| Naphthalene (NA) | <0.001 | <0.001 | <0.001 | 0.003 | 0.002 | 0.004 | 0.003 |
| Acenaphthene (AC) | <0.001 | <0.001 | <0.001 | 0.001 | 0.001 | 0.002 | 0.003 |
| Acenaphthylene (ACN) | <0.001 | <0.001 | <0.001 | 0.001 | 0.001 | 0.003 | 0.002 |
| Fluorene (FL) | <0.001 | <0.001 | <0.001 | <0.001 | 0.001 | 0.001 | 0.001 |
| Phenanthrene (PHE) | <0.001 | <0.001 | <0.001 | <0.001 | 0.001 | 0.001 | 0.001 |
| Anthracene (AN) | <0.001 | <0.001 | <0.001 | <0.001 | <0.001 | 0.001 | 0.003 |
| Fluoranthene (FA) | <0.001 | <0.001 | <0.001 | <0.001 | <0.001 | 0.001 | 0.002 |
| Pyrene (PY) | <0.001 | <0.001 | <0.001 | <0.001 | <0.001 | <0.001 | <0.001 |
| Benz[a]anthracene (B[a]A) | <0.001 | <0.001 | <0.001 | <0.001 | <0.001 | <0.001 | <0.001 |
| Chrysene (CHR) | <0.001 | <0.001 | <0.001 | <0.001 | <0.001 | <0.001 | <0.001 |
| Benzo[b]fluoranthene (B[b]F) | <0.001 | <0.001 | <0.001 | <0.001 | <0.001 | <0.001 | <0.001 |
| Benzo[k]fluoranthene (B[k]F) | <0.001 | <0.001 | <0.001 | <0.001 | <0.001 | <0.001 | <0.001 |
| Benzo[a]pyrene (B[a]P) | <0.001 | <0.001 | <0.001 | <0.001 | <0.001 | <0.001 | <0.001 |
| Indeno[1, 2, 3-cd]-pyrene (IP) | <0.001 | <0.001 | <0.001 | <0.001 | <0.001 | <0.001 | <0.001 |
| **Total PAH (mg/kg)** | <0.001 | <0.001 | <0.001 | **0.005** | **0.006** | **0.013** | **0.015** |

OG oil-polluted soil at day 1. GS good soil at day 1

Prominent bacterial isolates were *Bacillus* sp., *B. pumilis*, and *Pseudomonas aeruginosa* (Table 10) whereas*Mucormucedo*, and *Aspergillus niger*(Table 11) were the predominant fungi species during period of observation, irrespective of mushroom augmentation.

**Table 10:** Bacterial load of remediated soil at 2 months after inoculation

| Treatment | Bacteria load in cfu/g | Bacteria species |
| --- | --- | --- |
| Good Soil | $4.0 \times 10^4$ | *Staphylococcus* sp., *E. coli*, *Micrococcus varians*, *Pseudomonas aeruginosa*. |
| Control FO | $7.2 \times 10^4$ | *Proteus* sp., *P. vulgaris*, *B. pumilis*, *P. aeruginosa*, *Klebsiellasp*, *Bacillus* sp |
| 1 % FO | $6.0 \times 10^4$ | *Bacillus* sp., *B. pumilis*, *P. aeruginosa* |
| 5 % FO | $4.4 \times 10^4$ | *B. pumilis*, *P. aeruginosa*, *M. varians*, *Bacillus* sp |
| 10 % FO | $1.3 \times 10^4$ | *Proteus* sp., *Klebsiellasp*, *P. aeruginosa*, *B. pumilis*. |
| 25 % FO | $5.5 \times 10^4$ | *Azotobactersp.*, *M. varians*, *P. aeruginosa*, *Bacillus* sp. |
| 50 % FO | $7.0 \times 10^4$ | *E. coli*, *P. aeruginosa*, *Bacillus pumilis* |
| 100 % FO | $3.0 \times 10^4$ | *Azotobacter* sp., *M. varians*, *P. aeruginosa* |
| 1 % NO | $3.0 \times 10^6$ | *Bacillus* sp., *B. pumilis*, *P. aeruginosa* |
| 5 % NO | $1.8 \times 10^5$ | *Bacillus* sp., *B. pumilis*, *P. aeruginosa*, *M. varians* |
| 10 % NO | $1.5 \times 10^5$ | *Micrococcus* sp., *P. aeruginosa*, *B. pumilis*. |
| 25 % NO | $1.7 \times 10^4$ | *Azotobactersp.*, *M. varians*, *P. aeruginosa*, *B. pumilis*. |
| 50 % NO | $2.7 \times 10^4$ | *B. pumilis*, *P. aeruginosa*, *M. varians* |
| 100 % NO | $2.0 \times 10^6$ | *E. coli*, *P. aeruginosa*, *B. pumilis* |

**Table 11:** Fungi load (cfu/g) of polluted at soils 2 months after inoculation

| Treatment | Fungi load in cfu/g | Fungi species |
| --- | --- | --- |
| Good Soil | $9.0 \times 10^5$ | *Mucormucedo*, *A. niger*, |
| Control FO | $5.0 \times 10^4$ | *Mucormucedo*, *Aspergillus* sp., *Penicillum* sp. |
| 1 % FO | $6.0 \times 10^4$ | *Trichoderma hazianum*, *Penicilliumsp*, *Mucormucedo*, |
| 5 % FO | $2.8 \times 10^4$ | *MucorMucedo*, *Rhizopussp*, *Trichoderma hazianum*, *A. flavus*, |
| 10 % FO | $5.0 \times 10^4$ | *Mucormucedo*, *P. vulgaris* |
| 25 % FO | $4.4 \times 10^4$ | *Aspergillus* sp., *Rhizopussp*, *A. niger* |
| 50 % FO | $5.1 \times 10^4$ | *Mucormucedo*, *Trichoderma hazianum*, *A. flavus*, |
| 100 % FO | $6.0 \times 10^4$ | *Aspergillus* sp. *Rhizopus* sp. |
| 1 % NO | $1.2 \times 10^6$ | *Mucormucedo*, *Trichoderma hazianum*, *A. flavus*, *Penicilliumsp*. |
| 5 % NO | $3.0 \times 10^6$ | *MucorMucedo*, *Trichodermaharzianum* |
| 10 % NO | $1.0 \times 10^4$ | *Penicilliumsp*, *Aspergillus* sp., *Mucormucedo* |
| 25 % NO | $7.1 \times 10^4$ | *Rhizopussp*, *MucorMucedo* |
| 50 % NO | $3.3 \times 10^4$ | *MucorMucedo*, *Rhizopussp* |
| 100 % NO | $8.0 \times 10^6$ | *Aspergillus* sp., *Penicillum* sp., *Mucormucedo* |

## Discussion

The study investigated the use of a fungus, *Pleurotus tuberregium*in the remediation of petroleum polluted soils. The soils used were polluted severally to a chronic level at the point of collection, a situation that would have severely disturbed the soils. Several remediation strategies were utilized for the study, including, the exogenous introduction of mushrooms through sclerotia, supplementation using sawdust, provision of the enabling environment for the soil microorganisms and the soil seed bank. Crude oil pollution has been reported to impair free flow of air (oxygen) into the soil and also suppress the activities of microorganisms that would normally degrade the harmful substances (Basra *et al*., 2006).

The soils had a considerable number of bacteria and fungi species in the soils. *Bacillus*sp and *Klebsiella*sp occurred most among bacteria species and *Fusarium*spand*Aspergillus*sp among the fungi. Ogirri*et al*, (2001) observed that *Klebsiella*sp and *Aspergillus*sp were the prevalent species of bacteria and fungi respectively in oil polluted soils. Kwelang*et al*, (2008), reported that the fungal genera in polluted soils that they studied were Aspergillus, Penicillum and Mucor.

Following the harvest of mushrooms from the control and experimental soils, there wasgrowth of weeds in all the soils. These weeds sprouted from the soil seed bank. The weed species were identified in comparison with the weeds observed at the soil collection site at B-dere/Nabem communities. Anoliefo*et al*, (2006a, b) reported that the plant species that grew out of the experimental soils in their study were the same plant species observed within and around the auto mechanic workshops, where spent lubricating oil was constantly dumped.

Although remediation was obtained in the soil that not fungus-augmented, however, remediation was faster when soil was augmented with the fungus. Considerable reduction was observed in the present remediation study of Ogoni soils even when the termination date was still ahead. A remediation study does not necessarily have to be done for a prolonged time, rather, the right remediation strategies need to be employed. These strategies must include physical methods, involving periods of tillage and exposure of the sub and deep soils. There should be periods of application of water regimes and the use of supplementing and augmenting agents which would invariably boost the resident microorganisms. The use of mushrooms have been employed by many authors, however, the mushroom used for any specific study must be the type that are cultivated within and can easily adapt to the soil of the area of spill.

## References

Alexander, M. (2000) Aging, bioavailability and overestimation of risk from environmental pollutants. Environmental Science Technology 34: 4259–4265.

Anoliefo, G.O. and Ikhajiagbe, B. (2012) Techniques in Bioremediation. In (eds) Okhuoya, J.A., Okungbowa, F.Iand Shittu, H.O. Biological Techniques and applications; a comprehensive teaching manual. Uniben Press Benin City. pp. 211–232.

Anoliefo, G.O. and Umweni, F.O. (2004). Use of the edible African lettuce Launaetaraxcifolia to evaluate the remediation of soil contaminated with lead and cadmium. Nigerian Journal of Applied Sciences 22: 330–340.

Anoliefo, G.O., Isikhumhen, O.S., and Agbuna, S.O. (2001). Small-scale Industrial village in Benin-city, Nigeria: Establishment, failure and phytotoxicity assessment of soils from the abandoned site. Water, Air, and Soil Pollution Journal 131:169–183.

Anoliefo, G.O., Ikhajiagbe, B., and Dafe, F.V. (2006a) Ecotaxonomic distribution of plant species around motor mechanic workshops in Asaba and Benin City, Nigeria: Identification of oil tolerant plant species. African Journal of Biotechnology. 5(19): 1757–1762.

Anoliefo, G.O., Isikhuemhen, O.S., Eghwrudjakpor, K.O. and Iredia, H.O. (2006b). Growth of Chromolaenaodorata in crude oil polluted soil; an enhancement on the natural biodegradation process. Nigerian Journal of Botany 19(2): 360–376.

Anoliefo, G.O., Ikhajiagbe, B., Berena, A.T. and Okoro, R.E. (2010) Bioremediation of crude oil polluted soil by using Vernoniaamygdalina and manure. International Research Journal of Biotechnology 1(4): 037–043.

Barker, A.V., and Gretchen, M.B. (2004) Bioremediation of Heavy Metals and Organic Toxicants by Composting. The Scientific World Journal, 2:407–420.

Basra, S.M.A, Farooq, M, Afzal, I and Hussain, M. (2006) Influence of osmopriming on the germination and early seedling growth of coarse and fine rice. International Journal of Agricultural Biology. 8(1): 19–22.

Cal-EPA (2005). Air Toxics Hot Spots Program Risk Assessment Guidelines, Part II Technical Support Document for Describing Available Cancer Potency Factors. Office of Environmental Health Hazard Assessment, California Environmental Protection Agency. May 2005.

Dean, J. R. and Xiong, G. (2000). Extraction of organic pollutants from environmental matrices: selection of extraction technique. Trends in Analytical Chemistry, 19 (9): 553–564.

Efroymson, R.A., M.E. Will, and Suter, G.W. (1997a). Toxicological Benchmarks for Screening Contaminants of Potential Concern for Effects on Soil and Litter Invertebrates and Heterotrophic Process: 1997 Revision. Oak Ridge National Laboratory, Oak Ridge, TN. ES/ER/TM-126/R2 (http://www.hsrd. ornl.gov/ecorisk/reports.ht@.

Efroymson, R.A., M.E. Will, G.W. Suter, and Wooten, A. C. (1997b). Toxicological Benchmarks for Contaminants of Potential Concern for Effects on Terrestrial Plants: 1997 Retiision. Oak Ridge National Laboratory, Oak Ridge, TN, ES/ER/TM-85/R3 (http://www.hsrd.ornl.gov/ecoriskheports.html).

Esin (Eraydin) Erdogan and AytenKaraca (2011). Bioremediation of Crude Oil Polluted Soils. Asian Journal of Biotechnology, 3: 206–213.

Gunther, T., U. Dornberger and W. Fritsche, (1996). Effects of ryegrass on biodegradation of hydrocarbons in soil. Chemosphere, 33: 203–216.

Ikhajiagbe, B. (2010). Synergism in Bioremediation: Phytoassessment of Waste Engine Oil Polluted Soils after Amendment and Bioaugmentation. LAP Lambert Academic Publishing, Köln, Germany. 276p.

Ikhajiagbe, B and Anoliefo, G.O. (2012a). Substrate bioaugmentation of waste engine oil polluted soil. Research Journal of Environmental and Earth Sciences 4(1): 60–67.

Ikhajiagbe, B. and Anoliefo, G.O. (2012b). Phytoassessment of a 5-month old waste engine oil polluted soil after augmentation with Pleurotus tuberregium. Current Research Journal of Biological Science 4(1): 10–16.

Ikhajiagbe, B., Anoliefo, G.O., and Ogbogodo, J.B. (2015) Microbial quality and changes in polyaromatic hydrocarbon content of a bioremediated oil-polluted soil after exposure to different periods of tillage’ Nigerian Journal of Life Sciences, 5 (2): 90–97

Kalamees, R. and Zobel, M (2002) The role of seed bank in gap regeneration in calcacerous grassland community. Ecology 83: 1017–1025.

Nkwelang, G., Kamga, H.F.L., Nkeng, G.E., and Antai, S.P. (2008). Studies on the diversity, abundance and succession of hydrocarbon-utilizing microorganisms in tropical soil-polluted with oily sludge. African Journal of Biotechnology, 7: 1000–1080.

Ogirri, H.O., Ikenebomeh, M.J. and Anoliefo, G.O. (2001) Effect of oil pollution on the soil of an abandoned motomechanic village in Benin City. Tropical Science 41: 7–12.

Okhuoya JA, Isikhuemhen OS, Evue GA. 1998 –Pleurotus tuberregium(Fr.) Sing. sclerotia and sporophore yield during the cultivation on sawdust of different woody plants. International Journal of Mushroom Sciences 2, 41–46.

Parrish, Z.D., Banks, M.K. and Schwab, A.P. (2004) Effectiveness of phytoremediation as a secondary treatment for polycyclic aromatic hydrocarbons (PAHs) in composed soil. International Journal of Phytoremediation, 6(2): 119–137.

Plangklang, P. and Reungsang, A. (2008) Efects of rhizosphere remediation and bioaugmentation on caofuran removal from soil World Journal of Biotechnology 24: 983–989.

Udo, E.J., and Fayemi, A.A.A. (1975) The effect of oil pollution of soil on germination, growth and nutrient uptake of corn. Journal of Environmental Quality 4: 537–540.

